# Constitutive differences in glucocorticoid responsiveness are related to divergent spatial information processing abilities

**DOI:** 10.1101/579508

**Authors:** Damien Huzard, Avgoustinos Vouros, Silvia Monari, Simone Astori, Eleni Vasilaki, Carmen Sandi

## Abstract

The stress response facilitates survival through adaptation and is intimately related to cognitive processes. The Morris water maze task probes spatial learning and memory in rodents and glucocorticoids (i.e. corticosterone in rats) have been suggested to elicit a facilitating action on memory formation. Moreover, the early aging period (around 16-18 months of age) is susceptible to stress- and glucocorticoid-mediated acceleration of cognitive decline. In this study, we tested three lines of rats selectively bred according to their individual differences in corticosterone responsiveness to repeated stress exposure during juvenility. We investigated whether endogenous differences in glucocorticoid responses influenced spatial learning, long-term memory and reversal learning abilities in a Morris water maze task at early aging. Additionally, we assessed the quality of the different swimming strategies of the rats. Our results indicate that rats with differential corticosterone responsiveness exhibit similar spatial learning abilities but different long-term memory retention and reversal learning. Specifically, the high corticosterone responding line had a better long-term spatial memory, while the low corticosterone responding line was impaired for both long-term retention and reversal learning. Our modeling analysis of performance strategies revealed further important line-related differences. Therefore, our findings support the view that individuals with high corticosterone responsiveness would form stronger long-term memories to navigate in stressful environments. Conversely, individuals with low corticosterone responsiveness would be impaired at different phases of spatial learning and memory.

## INTRODUCTION

The stress response facilitates survival through physiological, behavioral and cognitive adaptations to the environment. It operates by facilitating organisms to cope properly with the demands of stressful situations (De Kloet et al., 2005; McEwen, 1998). The stress response is intimately related to cognitive processes. Glucocorticoid hormones - final products of the activated hypothalamus-pituitary-adrenocortical (HPA) axis-secreted during stress can impact learning and memory (Lupien et al., 1998; Sapolsky and Goosens, 2007). While chronic stress has detrimental effects on learning (Conrad, 2010), acute stress may induce either detrimental or beneficial cognitive changes, depending on the experimental and temporal context (de Quervain et al., 1998; Joëls et al., 2006; Lupien and McEwen, 1997; Salehi et al., 2010; Sandi and Pinelo-Nava, 2007; Schwabe and Wolf, 2010). For example, acute stress and acutely activated glucocorticoids delivered at synchrony with behavioral training can improve learning and memory (Sandi et al., 1997), whereas stress before a retention test can impair retrieval (de Quervain et al., 1998). It has also been shown that moderate stress can facilitate reversal learning (Graybeal et al., 2011).

In the Morris water maze - a task that probes spatial learning and memory in rodents (Morris, 1984) and depends on intact hippocampal function (Morris et al., 1990) - acute corticosterone (CORT) increases, associated with training, facilitate acquisition and subsequent retention of the task (Akirav et al., 2002; Conboy and Sandi, 2010; Sandi et al., 1997). This and other data (Cordero and Sandi, 1998; Oitzl et al., 1998; Roozendaal et al., 2006) have suggested a facilitating action for CORT on memory formation (for reviews, see de Quervain et al., 2009; Sandi, 2011). However, other studies have highlighted the existence of an inverted U-shape function for acute stress and glucocorticoid effects in memory formation (Luksys et al., 2009; Salehi et al., 2010; Sandi, 2013, 2011). Accordingly, whereas mild stress tends to facilitate memory processes, both low and high stress levels seem to be detrimental to memory function (Luksys and Sandi, 2011; Sandi, 2013, 2011). Typically, evidence for this view so far has included variations in stressor intensity (e.g., changes in the water temperature at which animals are tested in the water mazes). However, much less is known regarding the importance of individual differences in glucocorticoid activation for memory processes.

Our laboratory has performed selective breeding of rats according to their individual differences in CORT responsiveness to repeated stress exposure during juvenility (Walker et al., 2017). This selection resulted in high, intermediate and low CORT responders to stressful challenges (called ‘High’, ‘Inter’ and ‘Low’ lines, respectively). When tested during early adulthood, the progeny of these lines shows as well differences in stress responsiveness, with the opposite, High and Low, lines exhibiting high and low CORT levels, respectively, compared to the ‘normative’ Inter line (Huzard et al., 2019; Walker et al., 2017). In addition, these lines differ in other behavioral and autonomic nervous system responses (Huzard et al., 2019; Walker et al., 2017; Walker and Sandi, 2018). However, no previous information has been gathered for their performances in the cognitive domain.

Therefore, our goal here was to investigate whether endogenous differences in CORT responsiveness are associated with differences in spatial learning, long-term memory and reversal learning abilities in the water maze. Based on the role of CORT in the facilitation of memory acquisition during training and on the inverted U-shape theory, we hypothesized that the Low line would present worse spatial information processing capacities as compared to the Inter line. Moreover, as the CORT elevation displayed by the High line in response to a stress challenge (Walker et al., 2017) does not reach the high CORT levels found to induce cognitive impairments in a Morris water maze task (Salehi et al., 2010), we hypothesized that the High line would benefit from the endogenous higher glucocorticoids response and display superior spatial information processing capacities (particularly long-term memory) as compared to the Low and Inter CORT lines. To gain insight into the effects of stress on spatial learning abilities, we expanded the analysis of the traditional parameters from the Morris water maze by implementing a novel classification method that captures detailed aspects of swimming performances and exploratory strategies (Gehring et al., 2015; Vouros et al., 2018).

In addition, we remarked that there is currently no information linking different histories of endogenous glucocorticoid responsiveness to stress on different stages of spatial information processing. Given that the early aging period (around 16-18 months of age) is susceptible to reveal stress-induced (Bored et al., 2008; Sandi and Touyarot, 2006) and glucocorticoid-mediated (Bodnoff et al., 1995; Wheelan et al., 2018) acceleration of cognitive decline, we decided to test rats in the water maze during early aging. We avoided exposing animals to chronic stress, as it has been shown to accelerate cognitive decline during early senescence (Bodnoff et al., 1995; Bored et al., 2008; Sandi and Touyarot, 2006). Instead, we exposed animals to scattered behavioral challenges across life (juvenility, mid-age, and early aging) to assess whether differences in behavior and CORT responses in the selected lines are stable throughout life. This approach allowed us to interrogate whether our hypothesis regarding superior spatial information process abilities for High CORT responders would be evident during early aging.

## MATERIAL AND METHODS

### Animals

Experimental animals were 30 male Wistar Han rats from three rat lines selected for differential corticosterone reactivity (Huzard et al., 2019; Walker et al., 2017; see section *Protocol for selective breeding).* Animals were maintained on a 12:12 h light-dark cycle (lights ON at 07:00 h) in an environment controlled for temperature (22 ± 1 °C) and humidity (55 ± 5 % humidity). Rats had *ad-libitum* access to laboratory chow and water. All procedures were conducted in accordance with the Swiss National Institutional Guidelines on Animal Experimentation and were approved by the Swiss Cantonal Veterinary Office Committee for Animal Experimentation.

### Protocol for selective breeding

Rats used in this study were from the eighth generation (F8) of the corticosterone rat lines. They were mated with females at 5 months of age to generate F9 animals. The genetic selection process has been previously described (Huzard et al., 2019; Walker et al., 2017). Briefly, rats were selected following a ‘corticosterone-adaptation-stress-test’ (CAST) protocol that involved exposure to different stressors over three consecutive days during the juvenile period (P28-P30; ‘CAST’ on Figure 1). On P28, rats were exposed 5 min to an open arena followed by 25 mm on an elevated-platform (EP) in a bright room (> 300 lux). On P29, rats were placed in a new environment for 25 min where they were exposed to synthetic predator odor (trimethylthiazoline, TMT). Immediately afterwards, they were exposed during 25 min to an EP. On P30, the same stressors as on P29 were applied but in a reverse order. Two blood samples were taken from tail incision, on P28 and P30, one immediately following stress exposures and one alter 30 min of recovery in a neutral cage. Rats with high (> 250 ng/ml) or low (< 50 ng/ml) plasma CORT levels following stressor exposure on P30 were bred over generations, leading to the ‘High’ and ‘Low’ breeding lines, respectively. The control Inter line consisted of selectively bred animals with in-between CORT values following stressor exposure on P30.

**Figure 1.**
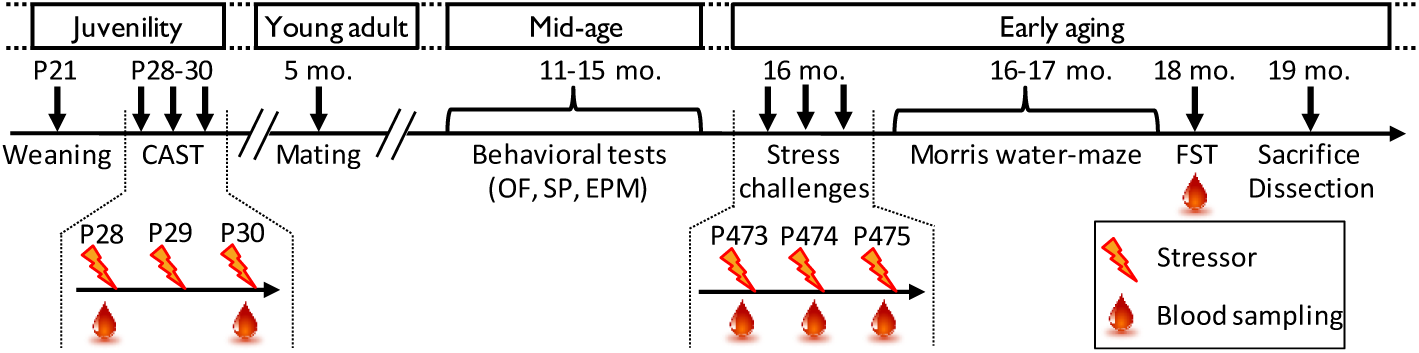
Outline of the experimental protocol. Rats from the eighth generation (F8) of CAST rats were weaned at postnatal day 21 (P21) and submitted to the CAST protocol between P28-30. At 5 months of age (5 mo.), young-adults were paired with females for one week in order to mate and generate the F9 generation of CAST rats. During the mid-age period, between 11 and 15 months of age (11-15 mo.), rats were tested on different behavioral tests (see Supplementary Material for details): an open field (OF), a social preference test (SP) and an elevated plus maze (EPM). Reaction to stress at early aging was assessed with 3 stress challenges applied at 16 months of age (16 mo.): rats were stressed with three different stressors over 3 consecutive days (P473-475) and blood was sampled following stress exposures. At 16 and 17 months of age (16-17 mo.) rats were trained in a Morris water maze task. At 18 months of age (18 mo.), rats were tested in a forced-swim test (FST) and blood was sampled. Finally, rats were sacrificed and adrenal glands weighted at 19 months of age (19 mo.). The orange and red lightning cartoon indicates a stress procedure. The red blood drop cartoon indicates blood sampling for determination of corticosterone in response to stress.

### Outline of the experimental protocol

This longitudinal study followed animals through different developmental periods, including i) pre-puberty (rats of one month of age were termed *juveniles*); ii) early adulthood (rats between 3 and 5 months of age rats were termed *young-adults*); iii) before reaching *mid-life* (around 12 months of age); and iv) the *early aging* period (between 15-19 months of age) (see Figure 1).

Rats were submitted to the CAST protocol at 1 month of age. Starting from 11 months of age, rats were exposed to behavioral challenges as described in Figure 1 and in Supplementary Methods. Reactivity to stress was tested at 16 months of age, when rats were submitted to a variety of stressors (bucket test, restraint stress and elevated platform) over three consecutive days. The Morris water maze was performed at 16-17 months of age. Rats were sacrificed and adrenal glands were dissected and weighted at 19 months of age.

### Responses to stress challenges during early aging

At 16 months ot age, rats were challenged with a different stressor per day over three consecutive days (Dl-3). On Dl, rats were placed for 30 min in an unescapable bucket (i.e. ‘bucket stress’). On D2, rats were submitted to 30 min restraint stress. On D3, rats were individually placed on an elevated platform in a bright room (> 300 lx) for 30 min. Immediately after exposure to each stressor, tail-blood was sampled for the analysis of CORT levels. In addition, on D2, blood was sampled at onset of stress, to assess basal CORT levels. These results are reported in the Supplementary Information file.

### Morris water maze

The maze consisted of a black circular pool (**∅** = 200 cm, 45 cm high) filled with 30 cm ot water at 23 ± 1 °C and virtually divided into four equivalent quadrants: northeast (NE), northwest (NW), southeast (SE) and southwest (SW). A circular rescue platform (**∅** = 12 cm; distance between platform center point and pool wall: 30 cm) was submerged 1-2 cm below the water surface. The testing room was illuminated (50 ± 10 lx) by lights placed below the pool to avoid light reflections. To monitor the animals, a camera was mounted to the ceiling above the center of the pool. The water maze was surrounded by extra-maze cues of different shape, size and color. A schematic sketch of the pool and of the experimental planning is represented on Supplementary Figure 1.

#### Spatial acquisition phase

The escape platform location (i.e. the *target*) remained the same for all trials of the acquisition phase (in the NE quadrant), whereas the starting location varied between trials in a semi-random design (East, South or West) as reported in Supplementary Figure 1. Rats were tested successively in blocks of four animals with an inter-trial interval of 5-10 min. Before starting the first training trial on the first day, each rat was placed tor 30 s on the platform. A trial began by placing the rat into the water facing the wall of the pool. The training trials were divided into 5 consecutive days (D1-5), with 5 trials on Dl-2 and 4 trials on D3-5. If a rat failed to escape within 120 s, it was guided by the experimenter to the target platform. After reaching and standing on the platform, a rat was left undisturbed for 15 s before being placed back in its homecage. For the data analysis, results From the two first days (Dl-2) and from the last two days (D4-5) of training were averaged.

#### Probe trial

On D5, 2 h after the last training trial, we assessed the reference memory with a retention probe trial during which the escape platform was removed from the pool and rats were allowed to swim tor 60 sec (D’Hooge and De Deyn, 2001; Vorhees and Williams, 2006). During the trial, rats started swimming From a new location (SW). In order to study the retention of the platform location, the latency to reach the target location and the time spent in the target (NE) quadrant were analyzed.

#### Long-term memory retention

On D17, in order to establish the long-term retention of the platform location, rats were tested on a second probe trial, after 12 days without training. This probe trial was followed by three retraining trials with the original platform location (target in NE quadrant).

#### Reversal learning

On D18, a reversal learning protocol was performed: the platform was moved to the opposite quadrant (target in SW quadrant) and rats were trained on four trials (starting positions: North, South, East and West respectively). Before the first reversal trial, rats were placed for 30s on the platform at the new location.

#### Analysis of water maze data

##### Classic water maze analysis

The mean latency to reach the platform was recorded. During each trial, animal’s movements were video-recorded and tracked by Ethovision software (Noldus, The Netherlands). The latency to reach the target, the time spent in the different quadrants, the velocity, the distance travelled before escaping and the cumulative distance to the target were analyzed.

##### Detailed classification of the swimming paths

We also performed an advanced analysis of the water maze data using the RODA software (Gehring et al., 2015; Vouros et al., 2018) which performs a detailed classification of the swimming paths into multiple strategies (See Supplementary Material). In brief this approach divided the trajectories into segments, which were classified into different classes of behavior. Results from this analysis led to a detailed categorization of swimming paths and allowed the detection of mixed strategies within a single trial (Gehring et al., 2015; Vouros et al., 2018). Eight different behavioral strategies were considered: Thigmotaxis, when an animal was swimming close to the walls of the arena; Incursion, when an animal started to move towards the inward locations of the arena; Scanning, when an animal was swimming randomly in the whole arena (Graziano et al., 2003); Focused Search, when an animal focused its search on a region of the arena different to the platform; Chaining Response, when an animal was swimming at the distance of the platform from the arena wall (Wolfer and Lipp, 2000); Sell Orienting, when an animal performed a loop while swimming, thereby orienting itself inside the arena (Graziano et al., 2003); Scanning Surroundings strategy, when an animal crossed the platform or the region around it; Scanning Target strategy when an animal performed a Focused Search within the platform region.

Some of the swimming strategies have been suggested to represent ‘low-level’ (or ‘suboptimal’) strategies (Janus, 2004; Vouros et al., 2018). During the Chaining Response strategy, an animal is not using the spatial cues but is instead finding the location of the platform with a non-spatial strategy, i.e. by having memorized the distance between the target and the pool’s wall (Broad and Holtzman, 2006; Janus, 2004; Wolfer and Lipp,2000). Other low-level swimming strategies are Thigmotaxis and Incursion. On the other hand. Self Orienting and Scanning Target strategies are considered as high-level strategies during which animals use the visual cues around the pool to orient themselves and find the platform (Gehring et al., 2015; Janus, 2004; Vouros et al., 2018).

### Forced-swim test

The CORT responses in response to 15 min of forced-swim test (FST) were assessed at 18 months ot age. Animals were individually placed in an unescapable plastic bucket (∅= 25 cm, 45 cm deep) containing 30 cm of water (23 ±1°C). Blood was sampled from the tail immediately after exposure, and blood plasma was extracted and analyzed to determine CORT concentration.

### Corticosterone analysis

Blood samples were collected into heparin-coated capillary tubes (Sarsted, Switzerland), kept on ice until centrifugation (4 min, 4°C and 9400 g), and stored at −20°C. Plasma CORT levels were measured using a highly sensitive ELISA kit (ADI-900-097, Enzo Life Sciences, Switzerland). Blood plasma samples were diluted 20 times and the ELISA was performed according to manufacturer’s instructions. Concentration values of CORT were calculated using a 4-parameter logistic fit (www.myassays.com). The intra- and inter-assay coefficients of variation were below 10%.

The percentage of CORT adaptation between P28 and P30 was computed with the formula: CORT adaptation (%) = ([CORT(P28)] - [CORT(P30)]) / [CORT(P28)] * 100.

### Statistics

Data distributions were controlled for normality and outliers were identified and removed by applying the Gaibbs’ (Grubbs, 1969) and ROUT (Motulsky and Brown, 2006) methods (with α= 5% and Q = 1 % respectively) from GraphPad Prism software (version 7.02). Two-way repeated measures ANOVAs, with *line* as a between-subject factor (three levels: High, Inter and Low lines) and time as a within-subject factor (number of levels depend on individual cases), were applied when data followed a repeated measures design. Other statistical analyses involved one-way ANOVA with *line* as a between-subject factor and *Post hoc* multiple comparisons were performed with a Fisher’s LSD test. When data did not follow a normal distribution, a non-parametric test (Kruskal-Wallis statistics followed by uncorrected Dunns’ multiple comparison) was applied. One rat from the High line was excluded from the experiment, in accordance with the Swiss Animal Experimentation Guidelines, due to critical health issue at 15 months of age (tumor growth on forelimb). Some blood samples were not obtained following the different stress challenges (P28,1 Inter and 2 High; P475,1 Inter; FST, 1 High). Videos from 2 Low line animals were not acquired during FST. Statistics and graphs were performed using GraphPad Prism. Statistical significance was set for at p < 0.05 for all tests. Data are presented as mean ± standard error of the mean (SEM).

## RESULTS

### Stable corticosterone and behavioral responses throughout life

So far, our work with these lines of rats selected tor their differential CORT responsiveness to stressors has verified that differential hormonal levels implied by the selection procedure at juvenility are still maintained when animals are exposed to life challenges during early adulthood (Walker et al., 2017). Here, we tested animals during the aging process. First, we verified that at P28 and P30 (CAST protocol; see Methods) there was a significant effect of line on CORT levels (Figure 2A: F_2,24_ = 40.22, p < 0.001). There was also an effect of time (F_2,24_= 15.55, p < 0.001) and a significant interaction between lines and time (F_2,24_ = 40.36, p < 0.001). Specifically, at P28 there was a difference in the amplitude of stress responses between the three lines with Low line having lower CORT levels compared to Inter (t_48_= 3.07, p = 0.004) and High lines (t_48_ = 2.97, p = 0.005). There was no difference in CORT levels between Inter and High lines at P28 (t_48_< 0.001, p = 0.99). At P30, Low line rats had significantly lower CORT values than both Inter (t_48_= 3,35 p = 0.002) and High (t_48_ = 1 1.95, p < 0.001) lines. At this time point, High line rats had as well a higher CORT response than the Inter line (t_48_ = 8.5, p < 0.001). Differences in CORT adaptation from P28 to P30 are illustrated in Figure 2B. There was a significant line effect on the percent change in CORT responses between P28and P30 (F_2,23_ = 128.4, p < 0.001). Low line animals habituated (i.e. positive adaptation) more to repeated stressor exposure than Inter (t_23_ = 2.87, p = 0.009) and High (t_23_ = 15.4, p < 0.001) lines, and the High line had a lower adaptation than the Inter line (t_23_= 11.9, p < 0.001). The Low and Inter lines had a lower CORT response at P30 than at P28, indicating habituation to stress (t-tests against 0: p < 0.001). On the opposite, High line rats exhibited a negative adaptation to stress (t-test against 0: p = 0.002) since they had a higher CORT response at P30 than at P28.

**Figure 2:**
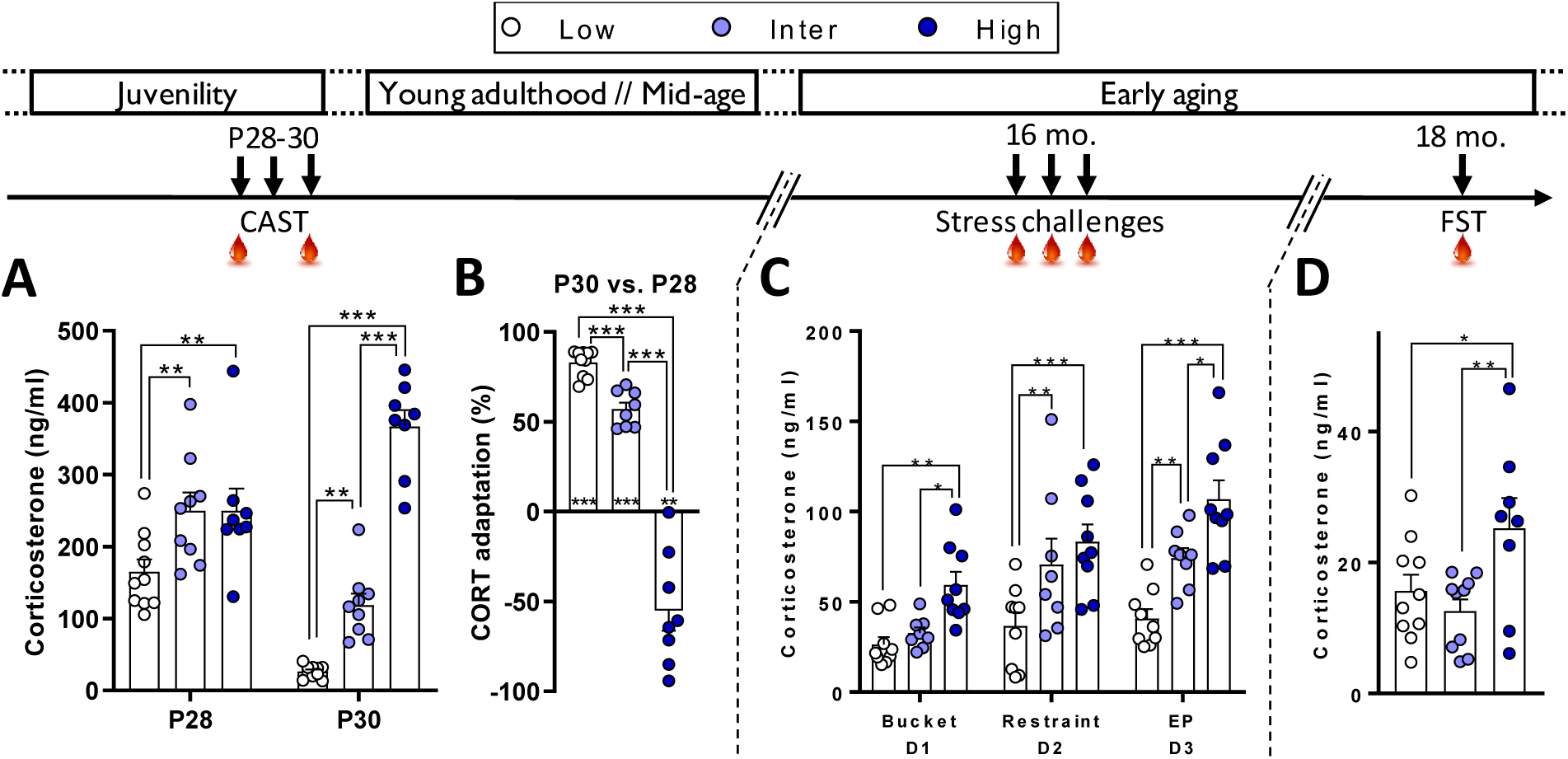
Different behavioral responses to stress between male rats from the CAST lines. **A**, At postnatal days 28 and 30 (P28 and P30), corticosterone (CORT) responses differed during CAST. At P28, Low line rats had a lower CORT response compared to Inter and High Lines. At P30, Low line rats had a lower CORT response in comparison to both Inter and High lines and High line animals had higher CORT compared to Inter line rats. **B**, The adaptation to stress, evaluated from the CORT response changes between P28 and P30, differed between the lines. Low line rats had a higher habituation (more positive adaptation) than Inter line rats and High line rats had a negative adaptation. Asterisks above columns indicate t-test statistics against 0% adaptation (i.e. no changes in CORT responses between P28 and P30). **C**, At 16 months of age (16 mo.), CORT responses after the exposure to three stressors showed the same line-dependent divergent profile as in younger rats. When CORT levels, following the three stressors, were analyzed independently, we observed that the Low line had lower CORT than the Inter line following exposure to restraint (p = 0.005) and EP (p = 0.005) but not after the bucket stress (p = 0.58). Moreover, Low line rats had lower CORT levels compared to the High line following the three stressors (p = 0.005, p < 0.001 and p < 0.001 respectively). There was a statistical difference in CORT response between Inter and High lines following the bucket stress (p = 0.025) and EP (p = 0.006) but not after restraint stress (p = 0.278). **D**, At 18 months of age (18 mo.), High line rats had higher CORT response to a forced-swim test (FST) than both Low and Inter lines. Low (n = 9-10), Inter (n = 9-10) and High (n = 8-9). Asterisks represent statistical differences: *p<0.05, **p<0.01, ***p<0.001.

Stress responsiveness was examined again at early aging (16 months of age) by exposing rats to a daily stressor (different one every day) over 3 consecutive days (Figure 2C). Again, CORT concentration in response to stress was different between the lines (F_2,23_= 15.1, p < 0.001). Low line animals had lower CORT responses than both Inter (t_23_= 2.71, p = 0.013) and High lines (t_23_= 5.5, p < 0.001). There was also a significant difference between Inter and High lines (t_23_= 2.63, p = 0.015).

At 18 months of age, we determined the CORT response following exposure to the forced-swim test (FST) (Figure 2D). There was a line effect on CORT values (F_2,25_= 4.67, p = 0.019). High line rats had higher CORT levels than both Low (t_25_= 2.25, p = 0.034) and Inter lines (t_25_= 2.97, p = 0.006), but there was no significant difference between Low and Inter lines (t_25_= 0.77, p = 0.45).

We also assessed the behavioral phenotype of the rats from the CAST lines at mid-age and early aging. There were no behavioral differences between the lines in an open field at 11 months of age (Supplementary Figure 2A). There was a statistical trend for a difference in sociability, at 14 months of age, with the High line being less social than the Low line (Supplementary Figure 2B). At 15 months of age, during an elevated plus maze, Low line rats were less anxious than both Inter and High lines (Supplementary Figure 2C). Finally, at 18 months of age, during the FST, High line rats did more passive floating than Inter line animals (Supplementary Figure 2D). At 19 months of age there were no differences in adrenal glands’ weight between the lines (Supplementary Figure 3).

### Spatial learning in the water maze

At 16-17 months of age, animals were tested for their spatial learning, retrieval, long-term memory and reversal abilities in the water maze. At training, animals demonstrated learning over subsequent training days, as indicated by their overall decrease in the latency to find the platform between D1-2 and D4-5 (F_1,25_ = 80, p < 0.001; Figure 3A). However, there were no significant differences in escape latencies between the lines (F_2,25_ = 2.17, p = 0.135), nor a significant interaction between lines and training days (F_2,25_ = 0.27, p = 0.77). However, there was a line effect for velocity of swimming (F_2,26_= 4.63, p = 0.019; Figure 3B); Inter line rats had higher swimming velocity than both Low (t_26_ = 2.22, p = 0.036) and High lines (t_26_ = 2.9, p = 0.007). These differences in velocity were visible at both D1-2 (Low vs. Inter, t_52_ = 2.11, p = 0.04; Inter vs. High, t_52_= 2.86, p = 0.006) and D4-5 (Low vs. Inter, t_52_= 1.78, p = 0.080; Inter vs. High, t_52_= 2.235, p = 0.030). The velocity of the rats during swimming decreased between D1-2 and D4-5 (F_1,26_ = 26.4, p < 0.001). There was no interaction betw’een line and training day (F_2,26_= 0.14, p = 0.81).

**Figure 3:**
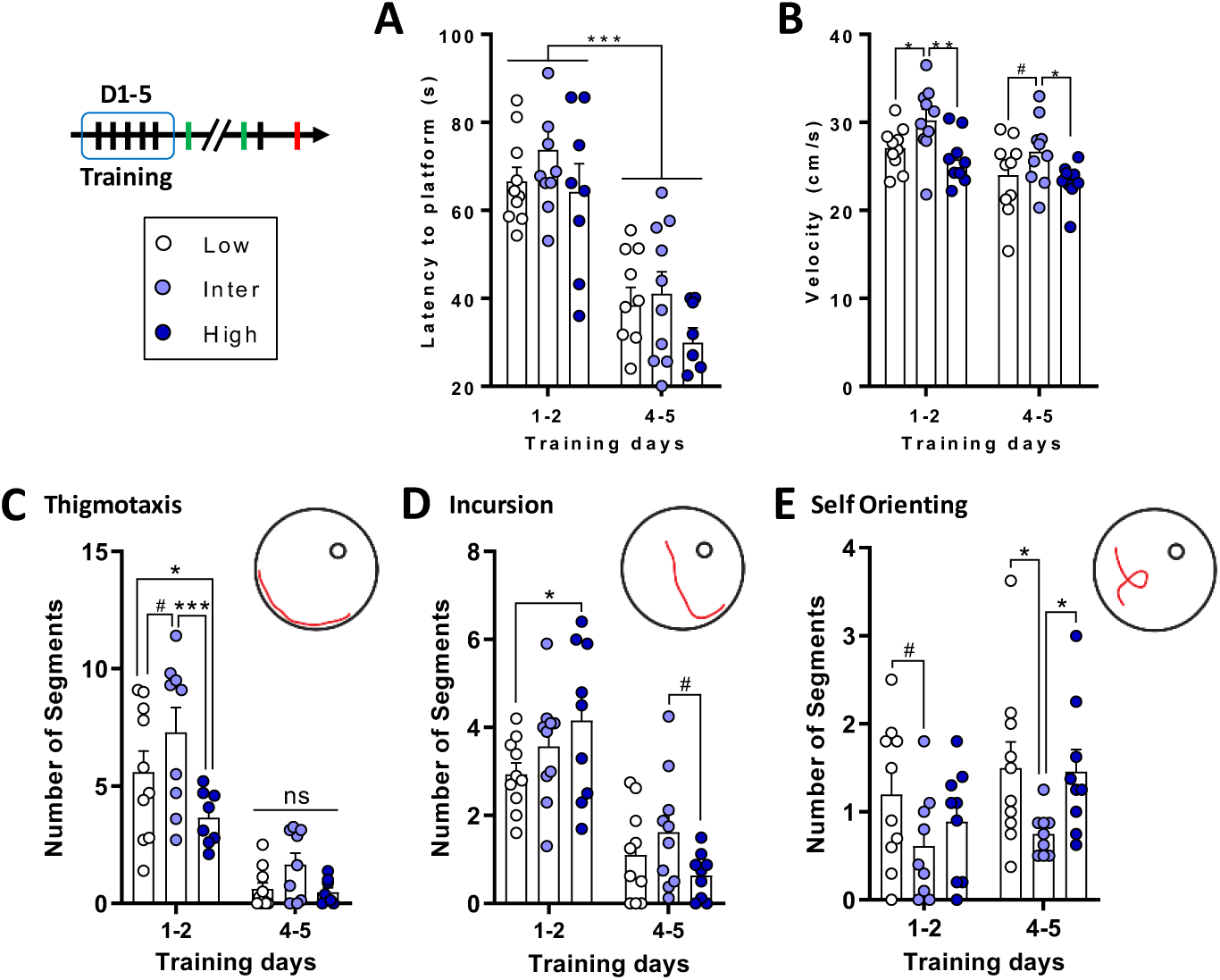
Similar escape latencies during water maze training between the lines, but inter line animals used ‘low-level’ swimming strategies. **A**, All rats showed similar learning and a decrease of the latency to escape the pool between D1-2 and D4-5. **B**, inter line rats had a higher swimming velocity than the Low and High lines. **C**, **D and E**, The different swimming strategies evaluated with RODA software (Vouros et al., 2018). **C**, Inter line rats used more the Thigmotaxis strategy compared to Low and High lines during D1-2 of water maze training. There were no differences in Thigmotaxis between the lines on D4-5 (ns). **D**, During D1-2 Low line rats did fewer incursions than High line rats. During D4-5, there was a tendency (# symbol) for a difference between inter and High lines. **E**, inter line rats used less Self Orienting on D4-5 than both Low and High line animals. Low (n = 10), inter (n = 9-10) and High (n = 8-9). Asterisks represent statistical differences: # p < 0.1, *p<0.05. **p<0.01. ***p<0.001. ns not significant.

An interesting picture emerged when swimming strategies were analyzed (see Figures 3C-E and Supplementary Figures 4D-H). For the Thigmotaxis strategy (Figure 3C), there was an overall effect of training days with a net decrease between D1-2 and D4-5 (F_1,24_= 77.7, p < 0.001). There was also a line effect (F_2,24_= 5.98, p An interesting picture emerged when swimming strategies were analyzed (see Figures 3C-E and Supplementary Figures 4D-H). For the Thigmotaxis strategy (Figure 3C), there was an overall effect of training days with a net decrease between Dl-2 and D4-5 (F_1.24_ = 77.7, p < 0.001). There was also a line effect (F_2.24_ = 5.98, p = 0.008); the Inter line used more this strategy than the Low (t_24_ = 2.06, p = 0.050) and High (t_24_ = 3.43, p = 0.002) lines, but there was no difference between Low and High line animals (t_24_ = 1.52, p = 0.141). There was also no interaction between lines and training days (F_2.24_ = 1.89, p = 0.173). For the Incursion strategy (Figure 3D), there was an overall effect of training days, with a decrease between Dl-2 and D4-5 (F_1,26_ = 73.6, p < 0.001). Although there was no line effect (F_2.26_ = 1.05, p = 0.364), we found a significant interaction between line and training day (F_2.26_ = 3.53, p = 0.044). At Dl-2, Low line rats did less Incursions than High lines rats (t_52_ = 2.24, p = 0.03); no differences were observed between the Inter line and both the Low (t_52_ = 1.19, p = 0.239) and High lines (t_52_ = 1.08, p = 0.286). No major differences were observed in this parameter at D4-5. For the Self Orienting strategy (Figure 3E), there was no training effect (F_1.25_ = 2.47, p = 0.128) or interaction between lines and training days (F_2.25_ = 0.33, p =0.72).

However, we found a significant line effect for this strategy (F_2.25_ = 6.32, p = 0.006). Inter line rats did less Self Orienting than both Low (t_25_ = 3.46, p = 0.002) and High lines (t_25_ = 2.48, p = 0.02), but no significant difference between Low and High lines (t_25_ = 0.91, p = 0.37). No major differences were found on distancetravelled (Supplementary Figure 4A, 4B), time spent in the target quadrant (Supplementary Figure 4G) or other swimming strategics (Supplementary Figures 4D-H and Supplementary Figure 5).

### Reference memory and long-term retention

Reference memory and long-term retention were tested through probe trials given on D5 (Figures 4A-C) and D17 (Figures 4D-G). On D5, there were no line-related differences in the time spent in the target quadrant (F_2,26_= 1.58, p = 0.225; Figure 4A), in the latency to reach the target (F_2,26_ = 0.487, p = 0.62; Figure 4B) and in swimming distance (F_2,24_= 1.7, p = 0.204; Figure 4C).

**Figure 4:**
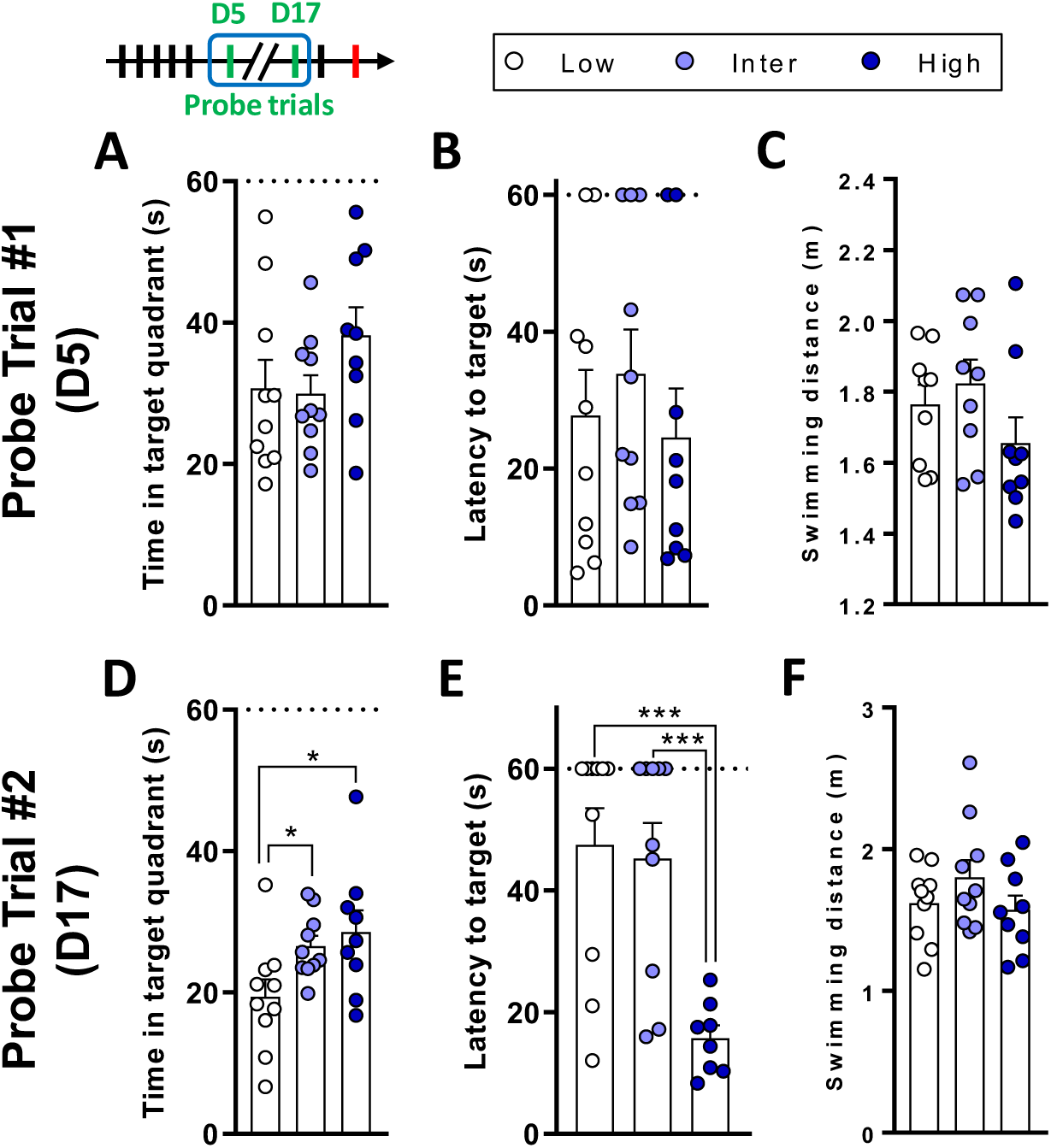
High line rats have better long-term spatial memory retention. **A. B and C**, Results from the first probe trial, performed on D5. **A**, The time spent in the target quadrant did not differ between the lines. **B**, All rats reached the target location with the same latency. **C**, There was no difference in the distance moved during the probe trial on D5. **D**, **E. and F**, Results from the second probe trial, performed on D17. **D**, Low line rats spent less time in the target quadrant compared to Inter and High lines. **E**, High line rats reached the platform location with a lower latency compared to Low and inter lines. **F**, There was no difference in the total swimming distance. Low (n = 9-10), inter (n = 9-10) and High (n = 8-9). Asterisks represent statistical differences: *p<0.05,***p<0.001.

On the long-term, second probe trial given on D17, there was a line effect on the time spent in the target quadrant (Figure 4D F_2.26_ = 4.16, p = 0.027). Low line rats spent less time in the target quadrant compared to Inter (t_26_ = 2.18, p = 0.038) and High line rats (t_26_ = 2.71, p = 0.012). Moreover, the latency to reach the target location (Figure 4E) differed between the lines (F_2,25_ = 10.3, p < 0.001). High line rats reached faster the target location than the Low (t_25_ = 4.13, p < 0.001) and Inter lines (t_25_= 3.84, p < 0.001). The distance moved (Figure 4F) did not differ between the lines (F_2,26_= 1.38, p = 0.27).

The analysis of the swimming strategies showed that on D17 Low line rats used less the Target Scanning strategy compared to the High line (see Supplementary Results and Supplementary Table 1). There were no additional differences in swimming strategies during the probe trials (Supplementary Table 1).

On D17, during the re-training trials with the target in the NE quadrant, there were no differences in the latency to reach the platform, in the distance traveled, in the swimming velocity, in the time spent in the target quadrant and in the average distance from the platform (Supplementary Figures 6A-E). Furthermore, there were no line-related differences in the swimming strategies (Supplementary Figure 5 and Supplementary Table 1).

### Reversal learning

Finally, on D18, we tested rats’ reversal learning abilities. To this end, the platform was placed in the opposite quadrant to previous trials. There was a line effect (F_2,26_=4.11,p=0.028) on the latency to reach the platform at the new location (Figure 5A). Low line rats had a longer latency to escape the water maze compared to Inter (t_26_ = 2.42, p = 0.023) and High line animals (t_26_= 2.52, p = 0.018). The distance traveled during reversal learning (Figure 5B) showed a tendency for a line effect (F_2,26_ = 2.61, p = 0.093). Low line rats traveled more distance than High line rats (t_26_ = 2.17, p = 0.039) and there was no difference between Inter line rats and both Low (t_26_= 1.67, p = 0.107) and High lines (t_26_= 0.548, p = 0.588).There were no differences on the velocity of swimming or on the time spent in the new target quadrant (Supplementary Figures 7A-B). Swimming strategies during reversal learning are represented in Figures 5C-E and reported in Supplementary Table I. For the Chaining Response strategy (Figure 5C). there was a significant line effect (Kruskal-Wallis H_3_= 6.22. p = 0.045). Inter line rats did more Chaining Responses than Low (Dunn’s p = 0.033) and High line rats (Dunn’s p = 0.030) and there was no difference between Low and High lines (Dunn’s p = 0.926). For the Tatgct Scanning strategy (Figure 5D), there was a significant line effect (Kruskal-Wallis H_3_= 6.37, p = 0.041). Inter line rats did less Target Scanning than Low line rats (Dunn’s p = 0.017) and there was a tendency for a difference between Inter and High lines (Dunn’s p = 0.061). There was no difference between Low and High lines in Target Scanning (Dunn’s p = 0.65). For the Self Orienting strategy (Figure 5E), there was a tendency for a line effect (F_2,26_=2.66. p = 0.089). Low line rats did more Self Orienting than High line rats (t_26_= 2.1. p = 0.045) and there was a statistical trend for a difference between Low and Interlines (t_26_ = 1.85, p =0.076).

**Figure 5:**
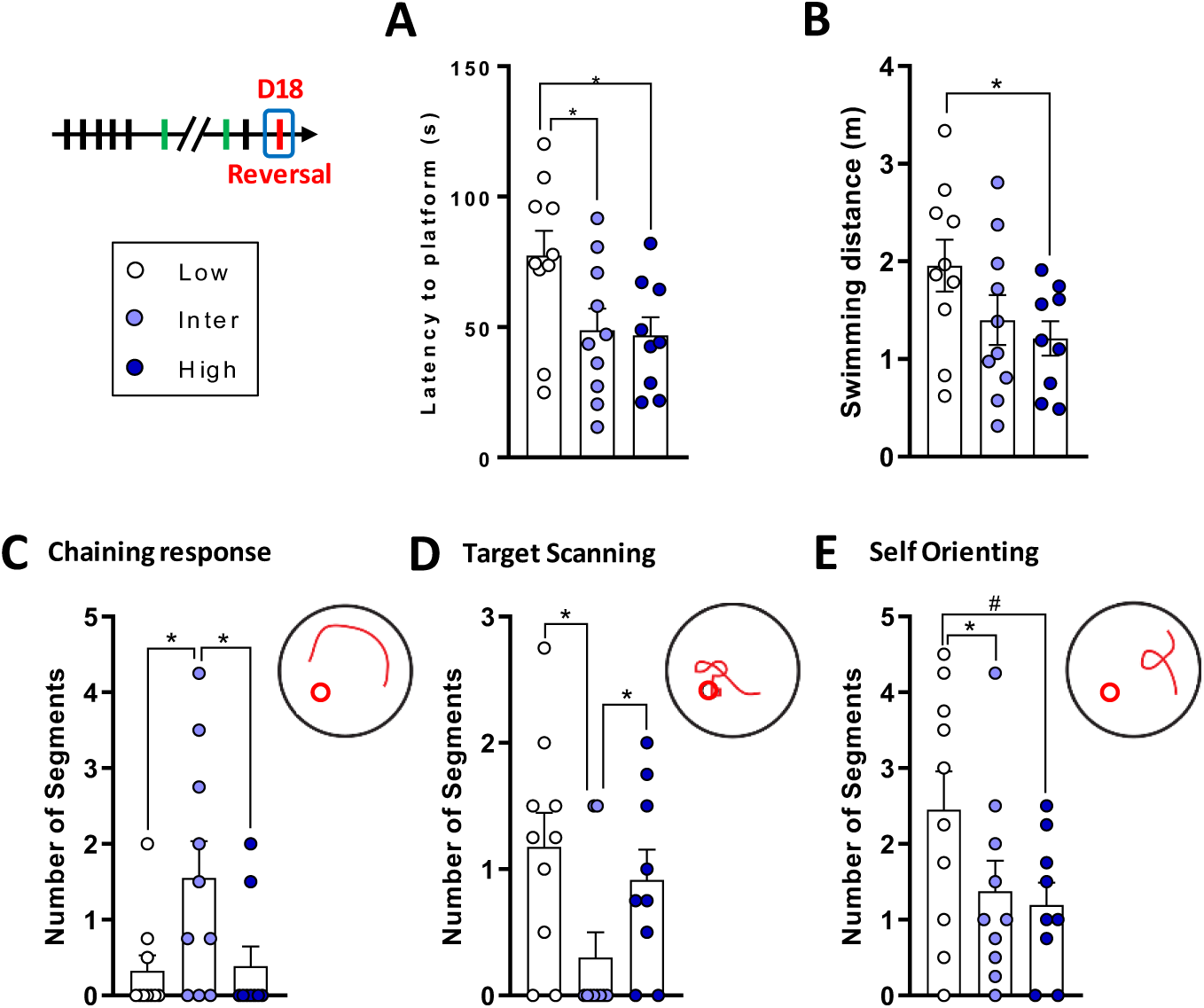
Low line rats show deficits in spatial reversal learning. **A**, Latency to find the platform in the new location. Low line rats spent more time to reach the target in comparison to Inter and High lines. **B**, There was a tendency for a difference on the distance traveled during reversal training with Low line rats swimming longer distances than High line animals. **C**, Inter line rats did more Chaining Responses compared to Low and High lines. **D**, Inter line rats did less Target Scanning than Low and High lines. **E**, Low line rats had a tendency to Self Orient more than Inter and High lines. Low (n = 10), Inter (n = 10) and High (n = 9). Asterisks represent statistical differences: # p < 0.1, *p<0.05.

There was no difference in Self Orienting between Inter and High lines (t_26_= 0.30, p = 0.76). There were no differences in the other swimming strategies during reversal learning (Supplementary Figure 5 and Supplementary Table 1).

## DISCUSSION

In the present study, w investigated spatial information processing abilities of rats that display constitutive differences in glucocorticoid responsiveness to stress. Our results indicate that rats with differential CORT responsiveness exhibit similar spatial learning abilities but different long-term memory retention and reversal learning. Specifically, the High CORT line had a better long-term spatial memory, while the Low line was impaired for both long-term retention and reversal learning. Interestingly, our analysis of performance strategies revealed important line-related differences.

Importantly, we confirmed that the three lines display a stable divergence in CORT responsiveness to stressors from the juvenile period throughout aging and a behavioral phenotype comparable to the one reported in young-adults from the same lines (Walker et al., 2017; Walker and Sandi, 2018; Huzard et al., 2019). Our results show that, as compared to Inter line rats, rats from the High line were high CORT responders, and those from the Low line were low CORT responders to stressful challenges throughout life.

In line with our hypothesis, High CORT line rats performed markedly better than the other two lines in the long-term memory test, whereas the Low line was the most disadvantaged. When considering the latency to reach the virtual platform during the long-term probe trial (taking place 12 days after the last training trial, on D17), there was a CORT line-graded effect: the High line showed shorter, the Inter line intermediate, and the Low line longer escape latencies.

Moreover, Low line rats spent a shorter time in the target quadrant. These findings align with previous studies showing that acute CORT secretion during learning of a spatial task improves long-term retention (Sandi et al., 1997; Akirav et al., 2004). Moreover, Low line rats were impaired in the reversal learning task. Given that their performance during the training phase was not different from the other two lines, the learning disadvantage during this second learning phase might be related to their impaired long-term memory of the task. The alternative possibility that the memory for the former platform location might have persevered interfering with the acquisition of the new platform location seems less plausible given that the Low line showed the lowest long-term memory retention in the D17 probe trial.

Previous studies showed that stress modulates the strategies used to navigate in spatial tasks in rodents and humans (Schwabe et al., 2007; Schwabe and Wolf, 2010; Schwabe et al., 2010, 2007; van Gerven et al., 2016) and that early life stressed rats tested at adulthood use more ‘low-level’ swimming strategies (Gehring et al., 2015; Vouros et al., 2018). Moreover, the stress response during a spatial task triggers a change in navigation strategies in order to protect against an impairment of performances (Schwabe et al., 2010; van Gerven et al., 2016). In order to rescue spatial performances, CORT may trigger a switch in memory systems leading to a change in the spatial strategy used (van Gerven et al., 2016). A mouse model of impaired water maze performance did not use spatial strategies during navigation but instead learned the location of the platform with a non-spatial strategy (i.e. Chaining strategy) which was defined as a use of ‘suboptimal’ (i.e. ‘low-level’) swimming strategies (Brody and Holtzman, 2006; Janus, 2004).

In our study, detailed classification of the swimming strategies revealed that across the different testing phases, rats from the Inter CORT line were the ones that systematically employed ‘low-level’ and fewer ‘high-level’ strategies (Gehring et al., 2015; Janus, 2004; Vouros et al., 2018) compared to Low and High lines. Thus, during training, they used more Thigmotaxis and, during reversal learning, they used more Chaining Responses and less Target Scanning and Self Orienting. We speculate that instead of using the visual cues around the pool (Self Orienting and Target Scanning), Inter line rats used the distance between the wall of the pool and the platform and swam fast and circularly (Thigmotaxis, Incursions and Chaining responses) to reach the target. Strikingly, High and Low CORT lines employed overall similar ‘high level’ strategies, consisting on high amounts of Self Orienting and Target Scanning, while low amounts of Chaining Response. However, there were important differences in the use of the strategies between these two rat lines that could contribute explaining the less advantageous performance of the Low line. Indeed, during the early training phase, the Low line displayed more Thigmotaxis than the High line, which instead performed more Incursions, a strategy that may provide better opportunities to learn spatial cues associated with the distance at which the platform is located. Moreover, more Self Orienting in the Low line during the reversal phase may have operated in detriment of the efficiency impinged by the Target Scanning strategy.

Based on the proposed inverted U-shape relationship between stress and spatial memory (Yau et al., 1995; Salehi et al., 2010; Sandi, 2011, 2013), impaired performance could have been expected for the High CORT line. However, our data indicates that CORT levels triggered by training in the High CORT line were most probably not abnormally high. When probed after the forced-swim test at 18 months of age, CORT responses across lines were lower than following stressful challenges CORT concentrations observed following stressor exposure during earlier life periods. In fact, an age-related decline in the adrenal stimulated secretion of corticosterone has been earlier documented (Cizza et al., 1994; Hauger et al., 1994; Oh et al., 2018). Moreover, it is important to note that the High CORT line shows normal CORT baseline and recovery levels following stress activation, while it displays higher peak CORT levels following stress exposure and lower habituation to repeated stress (Walker et al., 2017). Therefore, this pattern of mounting a high CORT response to stress while being capable to efficiently recover CORT levels to normal baseline values might have allowed the optimal cognitive performance of the High line during all cognitive phases.

Similarly, these glucocorticoid regulation characteristics might help as well explaining why the High CORT line did not show deleterious performance in the spatial task despite the fact that they were tested during early aging. This is particularly relevant given the substantial evidence in rodents and humans implicating midlife stress (Borcel et al., 2008; Sandi and Touyarot, 2006) and cumulative exposure to high glucocorticoid levels in age-related cognitive impairments (Bodnoff et al., 1995; Buechel et al., 2014; Landfield et al., 1981; Lupien et al., 1998; Lupien and McEwen, 1997; Wheelan et al., 2018; Yau et al., 2015). It was also shown, in various species, that aged individuals are impaired in spatial reversal learning (Eppinger et al., 2011; Lai et al., 1995; Mongillo et al., 2013).

Future studies should address the neurobiological mechanisms involved in the observed differences in spatial information processing reported here. Previously, several processes have been implicated in the facilitating effect of glucocorticoids in memory formation, including: endocannabinoid (Atsak et al., 2015) and cAMP-dependent protein kinase (Barsegyan et al., 2010) signaling, involved as well in the deleterious actions of glucocorticoids in memory retrieval (P. Atsak et al., 2012; Piray Atsak et al., 2012; Barsegyan et al., 2010); protein synthesis and synaptic glycoproteins (Bisaz et al., 2009; Merino et al., 2000; Sandi et al., 1995; Sandi and Rose, 1997); or the glucocorticoid receptor (GR)-mediated activation of the mitogen-activated protein kinase (MAPK) signaling pathway and the downstream regulated immediate-early gene Egrl (Revest et al., 2005) and subsequent activation of the pre-synaptic vesicle-associated phosphoprotein synapsin-Ia/Ib (Revest et al., 2010). Regarding work performed specifically on the Morris water maze task, synaptic translocation of the Alpha-amino-3-hydroxy-5-methyl-4-isoxazolepropionic acid receptor (AMPAR) GluA2 subunit was causally implicated in stress- and glucocorticoid-induced facilitation of spatial learning and memory (Conboy and Sandi, 2010) and synaptic efficacy (Martin et al., 2009). Therefore, it is plausible that superior performance in the High CORT line may involve AMPAR-dependent mechanisms and/or several of the other mechanisms mentioned above. Conversely, in agreement with the findings that stress affects reversal learning (Graybeal et al., 2011) and that glucocorticoids modulate mechanisms involved in reversal (Bryce and Howland, 2015; Myers et al., 2014; Raio et al., 2017), our data suggest that the Low CORT responding line may exhibit altered neural activity in key brain regions involved in flexible behaviors. In fact, our former analyses of these rats indicate that the Low line exhibits lower basal activity in the ventral orbitofrontal cortex (OFC), prefrontal cortex (mPFC) and hippocampus as compared to the High line (Walker and Sandi, 2018). Impaired behavioral flexibility during reversal learning in the water maze has been reported for both mice with mPFC damage and aged rats with altered neural encoding in the OFC (Latif-Hernandez et al., 2016; Schoenbaum, 2006). Furthermore, it was shown that hippocampal synaptic plasticity processes mediate spatial reversal learning in the water maze (Dong et al., 2013).

Therefore, our findings add important evidence to the view that high CORT responsiveness to stressful challenges may facilitate learning processes and memory storage. Conversely, we present strong evidence that extreme low CORT responsiveness may be deleterious for different cognitive functions. These findings were apparent even though animals were tested during early aging, a period when cumulative glucocorticoid levels tend to be deleterious for cognitive function. Our approach brings forward the concept that constitutive differences in glucocorticoid responsiveness are associated with differences in cognitive processing. In the future, it would be relevant to assess whether exposing these lines of rats to chronic stress at mid-life would have a different impact on their cognitive trajectories during the aging process. So far, the results reported here favor the hypothesis that having an efficient HPA axis, capable of mounting strong CORT responses while recovering swiftly a normal hormonal baseline, may be highly advantageous for optimal cognitive gain; at least when life history is not heavily charged in stressful challenges.

## Acknowledgements

We would like to thank Tanja Goodwin, Isabelle Guillot-de-Suduiraut and Jocelyn Grosse for valuable technical support during behavioral experiments. This project has been supported by grants from the European Union’s Seventh Framework Program for research, technological development and demonstration [grant agreementNo. 603016 (MATRICS)], Swiss National Science Foundation (NCCR SYNAPSY, grants No. 51NF40-158776 and 51NF40 - 185897] and intramural funding from the EPFL to CS. The funding sources had no additional role in study design, in the collection, analysis and interpretation of data, in the writing of the report or in the decision to submit the paper for publication. This paper reflects only the authors’ views and the European Union is not liable for any use that maybe made of the information contained therein.

## Author Contributions Statement

D.H. and C.S. conceived and designed the study. D.H. performed experiments, analyzed data and wrote the first version of the manuscript. A.V. and E.V. performed the classification of swimming strategies. S.M. provided technical support for behavioral testing. C.S. obtained funding and supervised the study. All authors gave inputs and approved the final version of the manuscript.

