## Supplementary for "Constitutive differences in glucocorticoid responsiveness are related to divergent spatial information processing abilities"

### Supplementary Methods:

#### Animals

In addition to the experimental animals described in the main text, other animals were used for task setting. Specifically, male Wistar Han rats of 5 weeks of age were purchased from a commercial breeder (Charles River, L’Arbresles, France) and were used as social stimuli in the social preference test.

#### Open field test

The open field (OF) test was performed to evaluate exploratory behavior in mid-aged animals, at 11 months of age. It was conducted in a circular open arena (40 cm high, Ø = 1 m) by placing the animal facing the wall and letting it explore freely the apparatus for 10 min. The OF was virtually divided in three parts for analysis: a central disk (Ø = 25 cm), an intermediate zone (annulus with Ø = 25-75 cm) and the remaining wall zone (annulus with Ø = 75 cm - 1 m). The total distance traveled and the time in the different zones were analyzed during the test using Ethovision software (Noldus, The Netherlands). The arena was cleaned with a 5% ethanol solution between animals.

#### Social preference test

The social preference (SP) test was performed to evaluate sociability at 14 months of age with a protocol adapted from a previous study using a three-chambered social test apparatus (Veenit et al., 2013). The test was carried out in a rectangular polycarbonate apparatus containing three chambers (a center 20 × 35 × 35 cm; a left and a right compartments 30 × 35 × 35 cm). Dividing walls had removable doors controlling access to the left and right chambers. Left and right compartments contained a central Plexiglas cylinder (Ø = 15 cm), transparent and with small holes, where either a social (unfamiliar juvenile rat, 5 weeks old) or a non-social stimulus (plastic bottle) was placed. The cylinder allows visual, auditory and olfactory communication. The juvenile rats were first habituated to the three-chambered apparatus by placing them individually in the cylinder during 10 min for 3 consecutive days before testing. On the testing day, an experimental rat was placed in the middle chamber and allowed to explore for 5 min, with the doors closed in order to block the access to the side chambers. After the habituation period, the unfamiliar juvenile was placed in one of the cylinders and the object on the other side. Then, both doors were removed, and the subject rat was allowed to explore the entire apparatus for 10 min. The session was video-recorded and the time spent sniffing each cylinder was scored offline. A rat was considered exploring/sniffing the object or the juvenile when the nose was close to a cylinder and oriented toward its content. The entire apparatus was cleaned with 5% ethanol solution between each subjects.

#### Elevated plus maze

To assess anxiety-like behaviors at 15 months of age, the elevated plus maze (EPM) was used (Pellow et al., 1985). The apparatus, made of black PVC, consisted of a plus-shaped elevated platform (50 cm above the floor) with two opposite open arms (50 × 10 cm) and two opposite closed arms (50 × 10 × 38 cm). The luminosity was set at 15-16 lux at the apex of the open arms and at 3-6 Lux in the closed arms. The rats were placed individually in the maze facing a closed arm and were allowed to explore the apparatus for 5 min. The maze was cleaned with 5% Ethanol between each animals. Videos were recorded from the ceiling and were analyzed with Ethovision (Noldus, The Netherlands). The distance traveled and the time spent in the open and closed arms of the maze were analyzed. Previous studies showed that there was an important reduction in the time spent in the open arms of the EPM between 12 and 18 months of age (Bessa et al., 2005) with aged male rats spending significantly less time in the open arms and center zone compared to young adults (Andrade et al., 2003). Moreover, it was shown that, in old rats, the time spent in closed arms is correlated with anxiety level (Boguszewski and Zagrodzka, 2002). Therefore, we compared the time spent in the closed arms, as a proxy for anxiety-like behaviors, between the lines at 15 months of age.

#### Forced-swim test

The behavioral and CORT responses were assessed in response to 15 min of forced-swim test (FST) at 18 months of age. Animals were individually placed in an unescapable plastic bucket (Ø = 25 cm, 45 cm deep) containing 30 cm of water (23 ± 1 °C). Blood was sampled from the tail immediately after, and blood plasma was extracted and analyzed to determine CORT concentration. In addition, videos were recorded and active (swimming) and passive (floating) coping strategies were subsequently analyzed.

#### Morris water maze protocol

Supplementary Figure 1 is a schematic representation of the Morris water maze (MWM) setup and protocol used.

#### Swimming strategies analysis

All additional swimming strategies extracted from the swimming paths during the different testing phase in the MWM protocol are reported in Supplementary Figure 5 and in Supplementary Table 1.

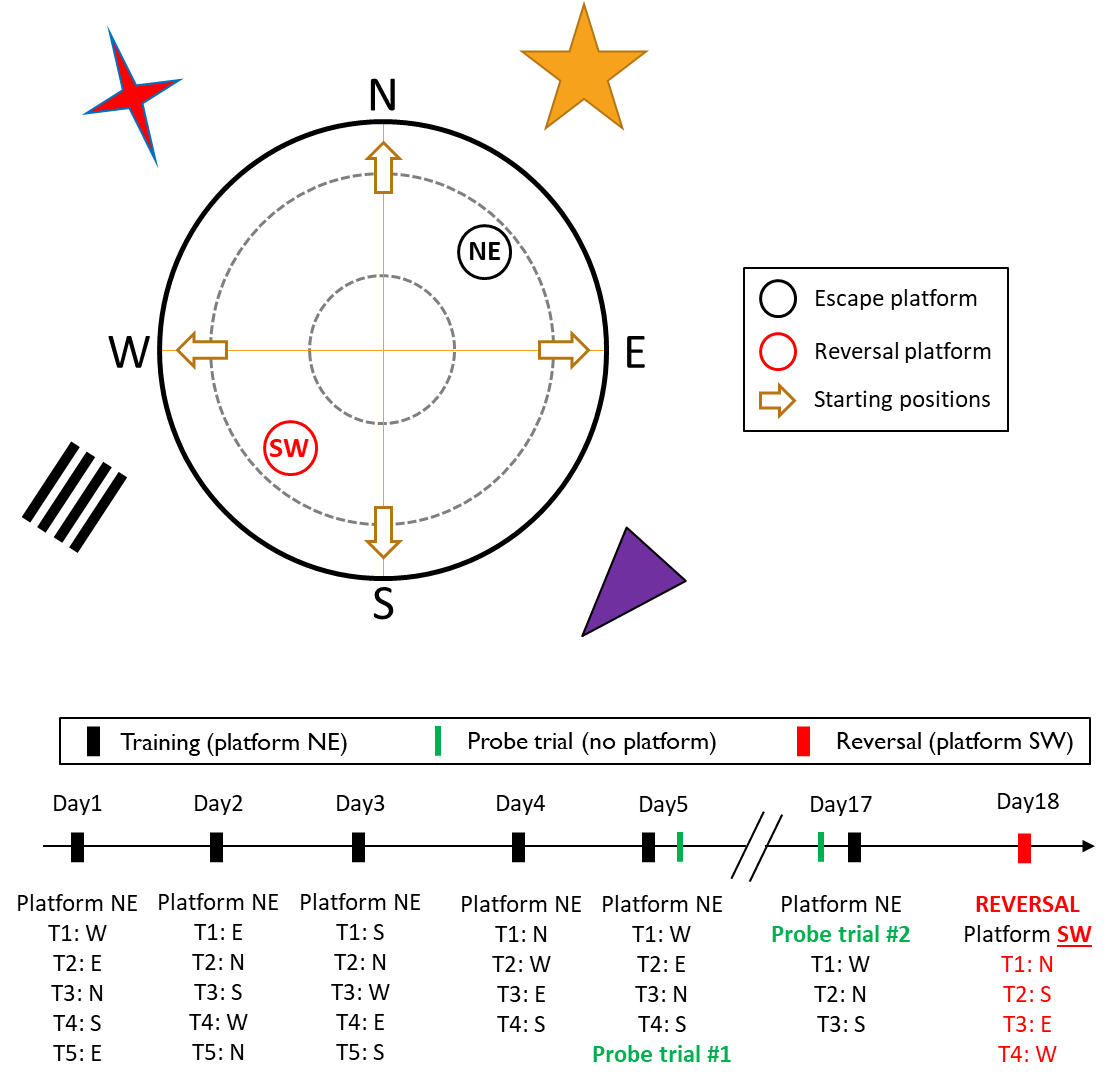

**Supplementary Figure 1. Schematic representation of the experimental protocol and planning of the Morris water maze task.** During day 1 to day 5 (D1-5), rats were training to escape the maze by reaching the platform in the northeast (NE) quadrant. On D5, a first probe trial was performed during 60 s after removal of the platform. On D17, a second probe trial was performed before 3 re-training trials. On D18, the platform was placed in the southwest (SW) quadrant and rats were trained for 4 trials. Starting positions followed a semi-random sequence and are indicated below each training days. The pool was surrounded by visual cues of different shape, size and color.

#### Adrenal glands weight

Rats were sacrificed, at 19 months of age, and both adrenal glands were dissected and weighed.

### Supplementary Results

#### Behavioral and emotional phenotypes and mid-age

We assessed the behavioral phenotype of the rats from the CAST lines at mid- and early aging. At 11 months of age, during the open field test (Supplementary Figure 2A), there was no difference in the amount of time spent in the central zone (F_2,26_ = 1.45, p = 0.25) and in the distance traveled (Supplementary Figure 2A, inner panel; F_2,25_ = 1.96, p = 0.16).

At 14 months of age, during the social preference test (Supplementary Figure 2B), there was a tendency for a difference in sociability index between the lines (F_2,27_ = 2.57, p = 0.095) and High line rats had a lower sociability than Low line animals (t_27_ = 2.25, p = 0.033). There were no differences between Inter line rats and both Low (t_27_ = 1.34, p = 0.192) and High lines (t_27_ = 0.91, p = 0.369).

At 15 months of age, during the EPM, there was a line effect (F_2,23_ = 6.36, p = 0.006) on the time spent in the closed arms (Supplementary Figure 2C). Low line rats spent less time in the closed arms of maze than both Inter (t_23_ = 2.49, p = 0.020) and High lines (t_23_ = 3.39, p = 0.003). There was no difference between Inter and High lines (t_23_ = 0.31, p = 0.306). There was no difference between the lines in the total distance traveled on the EPM (Supplementary Figure 2C, inner panel: F_2,23_ = 0.76, p = 0.48).

At 18 months of age, during the forced-swim test (Supplementary Figure 2D), there was a difference in floating time between the lines (F_2,23_ = 4.21, p = 0.028). High line rats spent more time floating than Inter line animals (t_23_ = 2.89, p = 0.008) but there was no difference with Low line rats (t_23_ = 1.36, p = 0.188).

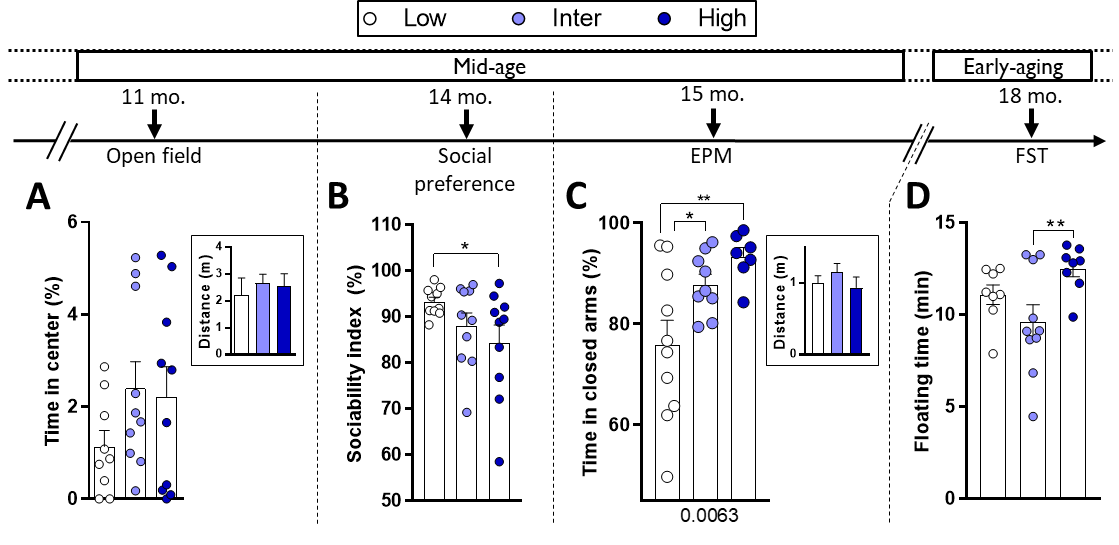

**Supplementary Figure 2. Differences in behavioral phenotypes of the lines throughout aging. A,** At 11 months of age (11 mo.), there was no difference in the time spent in the center of an open field (OF) and no difference in the distance traveled (**A, inset**). Low (n = 9), Inter (n = 10) and High (n =10). **B,** At 14 months of age (14 mo.), during a social preference (SP) test, there was a statistical tendency for difference in the sociability index between the three line (p = 0.095). High line rats had a lower sociability index than low line. Low (n = 9), Inter (n = 10) and High (n =10). **C,** At 15 months of age (15 mo.), there were differences in the time spent in the center and open arms of the elevated plus maze (EPM) between the three lines. Low line rats spent more time in the center and open arms compared to Inter and High lines. Low (n = 10), Inter (n = 9) and High (n =7). **D,** At 18 months of age (18 mo.), there was a line effect on the floating time during the forced-swim test (FST) and High line rats floated more than the Inter line rats. Low (n = 8), Inter (n = 10) and High (n =8). Asterisks represent statistical differences: *p<0.05, **p<0.01.

#### Adrenal glands weight

At 19 months of age, adrenals glands were extracted and weighed in order to establish long-term effects of stress on physiology (Blanchard et al., 1995). There was no difference in normalized adrenal glands’ weight between the lines (Supplementary Figure 3; p = 0.203).

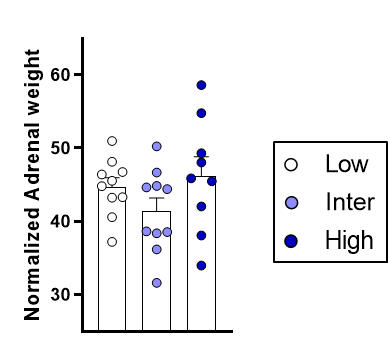

**Supplementary Figure 3. Adrenal glands’ weight.** Adrenal glands were weighed at 19 months of age and did not differ between the lines.

#### Supplementary MWM results

##### Training on D1-5

The distance traveled by the rats from the three lines before finding the platform (Supplementary Figure 4A) decreased between D1-2 and D4-5 (F_1,25_ = 117, p < 0.001). There was a statistical tendency for a difference in swimming distance between the lines (F_2,25_ = 2.77, p = 0.082) but there was no interaction between lines and days (F_2,25_ = 0.23, p = 0.79).

No line effects were observed in the following parameters: cumulative distance to the platform during swimming (Supplementary Figure 4B: Line, p = 0.243; Time, p < 0.001; Time × Line, p = 0.916); time spent in the NE platform (Supplementary Figure 4C: Line, p = 0.369; Line, p = 0.004; Time × Line, p = 0.377); Scanning behavior (Supplementary Figure 4D: Line, p = 0.169; Time, p < 0.001; Time × Line, p = 0.321); Focused search behavior (Supplementary Figure 4E: Line, p = 0.667; Time, p = 0.007; Time × Line, p = 0.502); Chaining Response behavior (Supplementary Figure 4F: Line, p = 0.538; Time, p = 0.003; Time × Line, p = 0.538); Scanning Around behavior (Supplementary Figure 4G: Line, p = 0.568; Time, p < 0.001; Time × Line, p = 0.158); and Target Scanning behavior (Supplementary Figure 4H: Line, p = 0.718; Time, p = 0.039; Time × Line, p = 0.919).

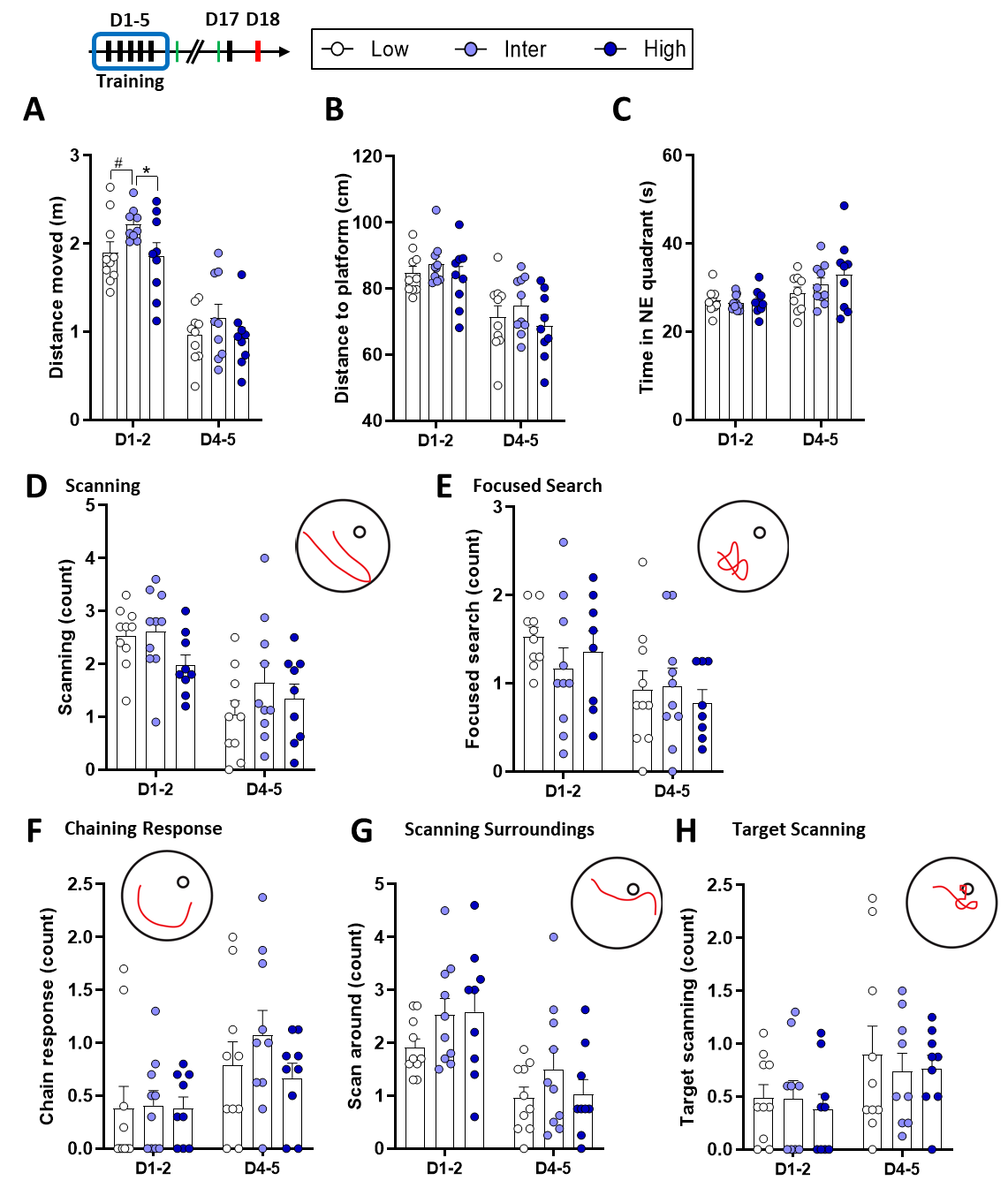

**Supplementary Figure 4: Additional results obtained during the D1-2 and D4-5 of MWM training with two-ways ANOVA summaries. A,** There was a statistical tendency for a line effect between the lines on the distance moved (p = 0.082). Low (n = 10), Inter (n = 9), High (n = 9); 1 Inter line rat was removed as outlier. **B,** There was no line effect on the cumulative distance to the platform during swimming (p = 0.243). **C,** There was no line effect on the time spent in the NE platform (p = 0.369). **D,** There was no line effect on the Scanning behavior (p = 0.109). **E,** There was no line effect on the Focused search behavior (p = 0.667). **F,** There was no line effect on the chain response behavior (p = 0.538). **G,** There was no line effect on the scanning around behavior (p = 0.158). **H,** There was no line effect on the target scanning behavior (p = 0.718). **B-H,** Low (n = 10), Inter (n = 10), High (n = 9).

##### Radar chart of the swimming strategies during MWM

Swimming strategies from all the training sessions were expressed as percentages and are represented in radar chart on Supplementary Figure 5.

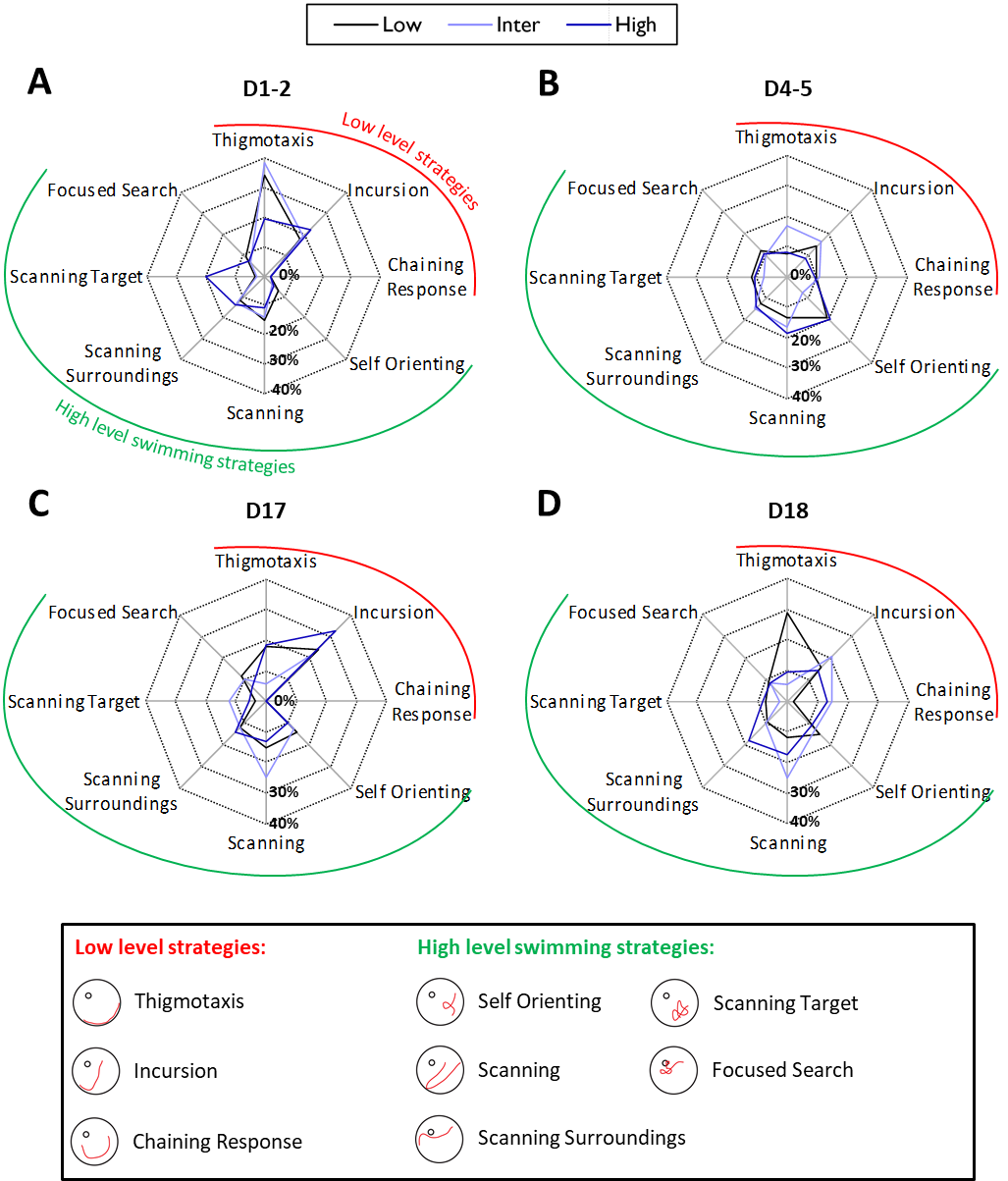

**Supplementary Figure 5: Radar charts of the percentage (%) of the different swimming strategies during MWM training. A,** During D1-2 of training, the High line did relatively more Scanning Target and less Thigmotaxis compared to both Low and Inter lines. **B,** during D4-5 Inter line did more Thigmotaxis and less Self Orienting. **C,** During re-training on D17, High line did more Incursion and Inter line did more Scanning strategy. **D,** During reversal learning on D18, Low line did more Thigmotaxis and less Chaining Responses, High line did more Scanning Surroundings and Inter line did more Incursions and Scanning strategies.

##### Probe trials on D5 and D17

During the two probe trials, Low line rats did less Target Scanning compared to High line animals on D17 (see Supplementary Table 1) but there were no additional differences between the lines in swimming strategies (see Supplementary Table 1).

Due to the low number (n = 2) of rats from the Low line displaying the Target Scanning strategy (see Supplementary Table 1) and due to the non-normality of the data distribution we used a non-parametric Kruskal-Wallis test. This statistical test was close to a statistical trend for a difference in Target Scanning during the probe trial of D17 (H_3_ = 4.428, p = 0.109) with a significant difference between Low and High line rats (Dunn’s p = 0.039). Additionally, we specifically tested whether Low line rats initiated less into Target Scanning strategy than High line rats with a chi-square test. The chi-square statistic (Χ^2^ = 4.2318) and corresponding p-value (p = 0.0397) suggested a statistical difference between the Low and High lines on the number of Target Scanning.

##### Re-training on D17

At D17, following the probe trial, rats were re-trained to reach the platform in the NE. The distance traveled during re-learning (Supplementary Figure 6B) showed no difference at D17. There was no difference in the swimming velocity of the rats (Supplementary Figure 6C). **D,** There was no difference in the time spent in the platform (NE) quadrant (Supplementary Figure 6D). **E,** There was no difference in the cumulative distance to the platform in the NE quadrant (PFNE) (Supplementary Figure 6E).

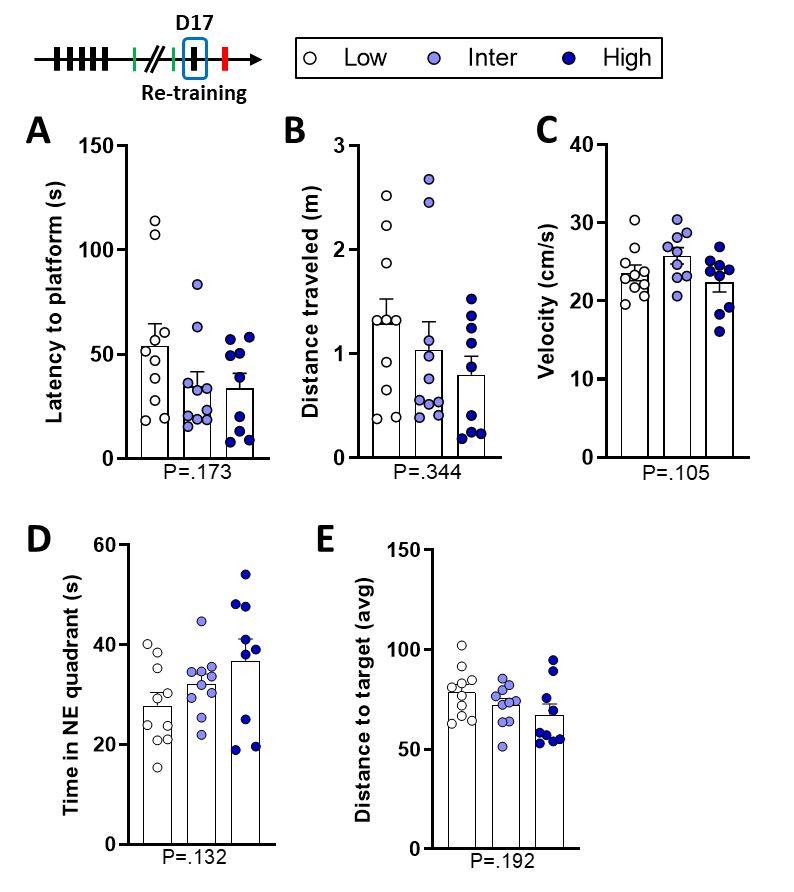

**Supplementary Figure 6. There were no differences during re-training on day 17. A,** There was no difference in the latency to reach the platform in the NE quadrant (p = 0.173). **B,** There was no difference in the distance traveled between the lines (p = 0.344). **C,** There was no difference in the swimming velocity of the rats (p = 0.105). Low (n = 10), Inter (n = 9), High (n = 9); 1 Inter line rat was excluded as outlier. **D,** There was no difference in the time spent in the platform (NE) quadrant (p = 0.132). **E,** There was no difference in the cumulative distance to the platform in the NE quadrant (PFNE) (p = 0.192). **A, B, D and E,** Low (n = 10), Inter (n = 10), High (n = 9). P-values of the one-way ANOVA are reported below each graph.

##### Reversal learning test on D18

For the Chaining Response strategy (Figure 5C), there was a significant line effect (Kruskal-Wallis H_3_ = 6.22, p = 0.045). Inter line rats used more the Chaining Response strategy than Low (Dunn’s p = 0.033) and High line rats (Dunn’s p = 0.030). There was no difference between Low and High lines (Dunn’s p = 0.926). A chi-square test was performed and described a similar difference between the lines (Χ^2^ = 6.7213, p = 0.0347).

There was no line effect (F_2,25_ = 2.01, p = 0.154) on the velocity of swimming during reversal trials (Supplementary Figure 7A).

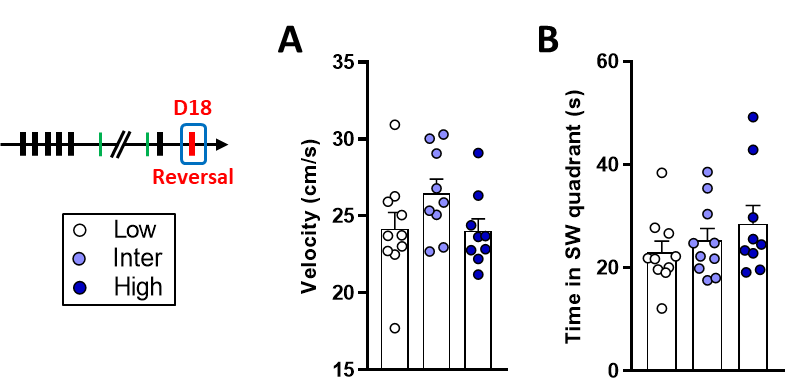

**Supplementary Figure 7. Extra-results from the reversal learning on day 18. A,** There was no difference in the velocity of swimming during reversal learning (p = 0.154). **B,** There was no difference in the time spent in the SW quadrant during reversal training (p = 0.355). Low (n = 10), Inter (n = 9-10), High (n = 9).

There was no difference in the time spent in the SW quadrant during reversal training (Supplementary Figure 7B) between the three lines (F_2,26_ = 1.08, p = 0.355).

| **Day** | **strategy** | **Low** | **Inter** | **High** | **one-way ANOVA** | | **Posthoc (or alternative statistics)** | **outliers** |
| --- | --- | --- | --- | --- | --- | --- | --- | --- |
| **Probe Trial #1 (D5)** | **Thigmotaxis** | 2.6 ± 0.98 (n=10) | 3 ± 1.29 (n=10) | 2.25 ± 0.62 (n=8) | F (2, 25) = 0.12 | P=.885 |  | 1 High |
|  | **Incursion** | 2 ± 0.61 (n=10) | 2.4 ± 0.62 (n=10) | 3.4 ± 0.87 (n=9) | F (2, 26) = 1.10 | P=.347 |  |  |
|  | **Scanning** | 2 ± 0.55 (n=9) | 3.4 ± 0.7 (n=10) | 2.8 ± 0.82 (n=9) | F (2, 25) = 1.02 | P=.375 |  | 1 Low |
|  | **Focused search** | 1.6 ± 0.78 (n=10) | 1.3 ± 0.54 (n=10) | 0.7 ± 0.44 (n=9) | F (2, 26) = 0.59 | P=.564 |  |  |
|  | **Chain response** | Not enough chain responses performed | | |  |  |  |  |
|  | **Self orienting** | 1.2 ± 0.59 (n=10) | 0.9 ± 0.48 (n=10) | 1 ± 0.71 (n=9) | F (2, 26) = 0.068 | P=.934 |  |  |
|  | **Scan around** | 3.1 ± 0.79 (n=10) | 3.6 ± 0.78 (n=10) | 1.7 ± 0.29 (n=9) | F (2, 26) = 2.09 | P=.144 |  |  |
|  | **Target scan** | 2.3 ± 0.92 (n=10) | 1.7 ± 0.65 (n=10) | 3.4 ± 0.99 (n=9) | F (2, 26) = 1.04 | P=.368 |  |  |
| **Probe Trial #2 (D17)** | **Thigmotaxis** | 2 ± 0.44 (n=9) | 3.4 ± 1.28 (n=10) | 1.12 ± 0.44 (n=8) | F (2, 24) = 1.66 | P=.212 |  | 1 Low, 1 High |
|  | **Incursion** | 2.2 ± 0.66 (n=10) | 3.1 ± 0.89 (n=10) | 2.1 ± 0.54 (n=9) | F (2, 26) = 0.58 | P=.567 |  |  |
|  | **Scanning** | 1.8 ± 0.55 (n=10) | 1.6 ± 0.41 (n=9) | 2.1 ± 0.7 (n=9) | F (2, 25) = 0.23 | P=.793 |  | 1 Inter |
|  | **Focused search** | 1.9 ± 0.59 (n=10) | 0.7 ± 0.37 (n=10) | 1.3 ± 0.6 (n=9) | F (2, 26) = 1.36 | P=.275 |  |  |
|  | **Chain response** | Not enough chain responses performed | | |  |  |  |  |
|  | **Self orienting** | 1.5 ± 0.37 (n=10) | 2 ± 0.79 (n=10) | 1.2 ± 0.6 (n=9) | F (2, 26) = 0.41 | P=.667 |  |  |
|  | **Scan around** | 3.4 ± 1.07 (n=10) | 2.4 ± 0.65 (n=10) | 3.2 ± 0.91 (n=9) | F (2, 26) = 0.37 | P=.697 |  |  |
|  | **Target scan** | 0.4 ± 0.27 (n=10) | 1.3 ± 0.54 (n=10) | 1.8 ± 0.55 (n=9) | KW H(3) = 4.43 | P=.109 | **Non-parametric: Low < High p = 0.039**  Chi-square Low vs High: Χ^2^ = 4.23, p = 0.04 | Only 2 Low scanned target |
| **re-training (D17)** | **Thigmotaxis** | 1.4 ± 0.57 (n=9) | 0.33 ± 0.14 (n=8) | 1.1 ± 0.57 (n=9) | F (2, 23) = 1.23 | P=.310 |  | 1 Low, 2 Inter |
|  | **Incursion** | 1.9 ± 0.61 (n=10) | 1.2 ± 0.42 (n=10) | 1.9 ± 0.72 (n=9) | F (2, 26) = 0.43 | P=.655 |  |  |
|  | **Scanning** | 1.2 ± 0.25 (n=9) | 1.4 ± 0.32 (n=10) | 0.8 ± 0.3 (n=9) | F (2, 25) = 1.16 | P=.329 |  | 1 Low |
|  | **Focused search** | 0.9 ± 0.31 (n=10) | 0.58 ± 0.13 (n=10) | 0.4 ± 0.17 (n=9) | F (2, 26) = 1.49 | P=.244 | SQRT Transformation |  |
|  | **Chain response** | Not enough chain responses performed | | |  |  |  |  |
|  | **Self orienting** | 1.15 ± 0.29 (n=9) | 0.73 ± 0.2 (n=10) | 0.59 ± 0.21 (n=9) | F (2, 25) = 1.42 | P=.260 |  | 1 Low |
|  | **Scan around** | 0.97 ± 0.23 (n=10) | 0.7 ± 0.22 (n=10) | 0.85 ± 0.25 (n=9) | F (2, 26) = 0.34 | P=.717 |  |  |
|  | **Target scan** | 0.33 ± 0.12 (n=9) | 0.7 ± 0.24 (n=10) | 0.33 ± 0.17 (n=8) | F (2, 24) = 1.29 | P=.294 |  | 1 Low, 1 High |
| **Reversal (D18)** | **Thigmotaxis** | 4.7 ± 2.01 (n=10) | 0.56 ± 0.15 (n=8) | 1.06 ± 0.39 (n=9) | F (2, 26) = 1.25 | P=.53 | **Non-parametric test** |  |
|  | **Incursion** | 2.55 ± 0.5 (n=10) | 2.22 ± 0.64 (n=10) | 1.5 ± 0.45 (n=9) | F (2, 26) = 0.90 | P=.419 |  |  |
|  | **Scanning** | 1.92 ± 0.50 (n=10) | 2.65 ± 0.5 (n=10) | 1.9 ± 0.35 (n=9) | F (2, 26) = 0.87 | P=.431 |  |  |
|  | **Focused search** | 1.42 ± 0.21 (n=10) | 0.88 ± 0.25 (n=10) | 0.9 ± 0.21 (n=9) | F (2, 26) = 1.84 | P=.179 | SQRT Transformation |  |
|  | **Chain response** | 0.32 ± 0.20 (n=10) | 1.55 ± 0.49 (n=10) | 0.39 ± 0.26 (n=9) | KW H(3) = 6.22 | **P=.045** | **KW** / Χ^2^ Inter > Low / high (p=0.07 / 0.037) |  |
|  | **Self orienting** | 2.45 ± 0.50 (n=10) | 1.37 ± 0.40 (n=10) | 1.2 ± 0.3 (n=9) | F (2, 26) = 2.66 | ***P=.089*** | Low > Inter / High (p=0.076 / 0.045) |  |
|  | **Scan around** | 1.6 ± 0.36 (n=10) | 1.12 ± 0.32 (n=10) | 2.03 ± 0.56 (n=9) | F (2, 26) = 1.16 | P=.328 |  |  |
|  | **Target scan** | 1.17 ± 0.27 (n=10) | 0.3 ± 0.2 (n=10) | 0.92 ± 0.24 (n=9) | KW H(3) = 6.37 | **P=.041** | **KW: Inter < Low / High (p = 0.017 / 0.061)** | Only 2 Inter scanned target. |
| **Supplementary Table 1. Mean ± SEM and statistical analysis of swimming strategies during probe trials #1 and #2, re-training and reversal training.** One-way ANOVA was computed for the different swimming strategies when data were normally distributed. In case data were not normally distributed, non-parametric Kruskal-Wallis and Chi-squared statistics were computed (in red font). The last column indicates the number of potential outliers excluded in order to respect normality of the data distributions. | | | | | | | | |
